# The structure of SALM5 suggests a dimeric assembly for the presynaptic RPTP ligand recognition

**DOI:** 10.1101/225821

**Authors:** Sudeep Karki, Prodeep Paudel, Celeste Sele, Alexander V. Shkumatov, Tommi Kajander

## Abstract

Synaptic adhesion molecules play a crucial role in the regulation of synapse development and maintenance. Recently several families of leucine rich repeat domain containing neuronal adhesion molecules have been characterized, including netrin G-ligands, LRRTMs, and the SALM family proteins. Most of these are expressed at the excitatory glutamatergic synapses, and dysfunctions of these genes are genetically linked with cognitive disorders, such as autism spectrum disorders and schizophrenia. The SALM family proteins SALM3 and SALM5, similar to SLITRKs, have been shown to bind to the presynaptic receptor protein tyrosine phosphatase (RPTP) family ligands. Here we present the 3 Å crystal structure of the SALM5 LRR-Ig domain construct, and biophysical studies that verify the crystallographic results. We show that both SALM3 and SALM5 extracellular domains form similar dimeric structures, in which the LRR domains form the dimer interface. Both proteins bind to the RPTP lg-domains with micromolar affinity. SALM3 shows a clear preference for RPTP-ligands with the meB splice insert. This is in accordance with previous results showing that the LRR domain is also required for the ligand binding. Our structural studies and sequence conservation analysis suggests a ligand binding site and mechanism for RPTP binding via the dimeric LRR domain region.

## 1. Introduction

Synaptic adhesion molecules play important role in the regulation of synapse development and maintenance, including the formation of early synapses and their differentiation into mature synapses (Missler *et al*., 2012; Südhof, 2008). These molecules are present on either presynaptic or postsynaptic side of the synaptic cleft on the neuronal cell membrane and contribute to the regulation of synapse development through trans-synaptic adhesion, either via homophilic interactions, as in the case of e.g. the NCAMs (Kallapur and Akeson, 1992) or heterophilic interactions, such in the case of the e.g. the neurexin and neuroligin (Sudhof, 2017). A large number of leucine rich repeat (LRR)-containing synaptic adhesion molecules such as the LRRTMs (Lauren *et al*., 2003), netrin-G ligands (NGLs) (Woo *et al*., 2009), synaptic adhesion-like molecules (SALMs) (Nam *et al*., 2011) and SLITRKs (de Wit and Ghosh, 2014) have been identified to be involved in synapse formation and maintenance, in particular in the excitatory synapses of the brain.

The SALM family of proteins consists of five members (SALM1-5) (Nam *et al*., 2011). They share a similar domain structure, containing seven LRRs, an immunoglobulin (Ig) domain and a fibronectin type III (Fn) domain in the extracellular region, a single transmembrane helix, and a cytoplasmic tail. In particular, subsets of SALMs (SALM1-3) possess a C-terminal type-I PDZ binding motif that can bind to PDZ domains of postsynaptic density protein PSD-95. All five SALMs are mainly expressed in brain, and have been implicated in the regulation of synapse development and function. SALM1 has been suggested to interact with other SALM proteins, and to interact with GluN1 subunit of NMDA receptors (NMDARs). SALM1 has further been reported to promote dendritic clustering of NMDARs in cultured neurons (Wang *et al*., 2006). SALM2 associates with both NMDARs and AMPA receptors (AMPARs), and promotes the development of excitatory synapses (Ko *et al*., 2006). SALM3 and SALM5 specifically induce both excitatory and inhibitory presynaptic differentiation in contacting axons via trans-synaptic interactions with type IIa presynaptic leukocyte antigen related (LAR)-family receptor protein tyrosine phosphatases (RPTPs) (Choi *et al*., 2016; Li *et al*., 2015).

The LAR-RPTPs consist of three members in vertebrates: LAR, RPTPσ, and RPTPδ, and each member contains three immunoglobulin (Ig) and four to eight fibronectin type III repeats and have multiple splice variants with short inserts at mini exon A, B and C sites (meA, meB and meC) in the extracellular region, with meA and meB in the Ig2 domain and between the Ig3 and Ig3 domains, respectively, and possess two tandem intracellular protein tyrosine phosphatase domains (see e.g. Takahashi and Craig, 2013, Um and Ko, 2013). LAR-RPTPs form multiple trans-synaptic adhesion complexes with various postsynaptic binding partners, including NGL-3 (Woo *et al*., 2009), SALM3 and SALM5 (Choi *et al*., 2016; Li *et al*., 2015) neurotrophic receptor tropomyosin-related kinase C (TrkC) (Takahashi *et al*., 2011), interleukin-1-receptor accessory protein-like1 (IL1RAPL1) (Yamagata *et al*., 2015), interleukin-1 receptor accessory protein (IL1RAcP) and SLITRKs (1–6) (Takahashi *et al*., 2012; Um and Ko, 2013; Yim *et al*., 2013). Thus, they appear to act as presynaptic hubs for synaptic adhesion and signaling. They can also act as signaling co-receptors for glypicans (Um and Ko, 2013).

SALM3 and SALM5 interact with the Ig-domains of all three members of LAR-RPTP family and have been reported to promote both the excitatory and inhibitory synapse development (Choi *et al*., 2016; Li *et al*., 2015). Intriguingly, SALM3 binds RPTP splice variants with insert at the meB site (Li *et al*., 2015), whereas SALM5 appears to bind to LAR-RPTPs independent of alternative splicing events at the meB site (Choi *et al*., 2016). A recent study showed that SALM4 suppresses excitatory synapse development by cis-inhibiting trans-synaptic adhesion interaction be SALM3 and LAR-RPTP (Lie *et al*., 2016). Further, SALM1 and SALM5 have been implicated in severe progressive autism and familial schizophrenia (Choi *et al*., 2016; Morimura *et al*., 2017).

The lack of a high-resolution structure for the SALM synaptic adhesion proteins has limited the understanding of how SALMs act in synapse formation and maintenance. Investigating the detailed molecular mechanisms underlying the SALM synaptic adhesion and other functions will be crucial for the further analysis of the biological function of these proteins. In the current study, we report the crystal structure of SALM5 extracellular LRR-Ig domain fragment, together with biophysical characterization of the protein family and binding to presynaptic ligand RPTPσ and its meB splice insert variant. Based on this we suggest a molecular model for the oligomerization and function of the SALM proteins in synaptic adhesion.

## 2. Methods

### 2.1. Cloning, expression and purification SALM and RPTP protein constructs

The mouse SALM gene constructs (SALM1 LRR-Ig-Fn_20-518_, SALM3 LRR-Ig_17-367_, SALM3 LRR-Ig-Fn_17-510_, and SALM5 LRR-Ig_18-376_ and SALM5 LRR-Ig-Fn_18-505_ and mouse RPTPσ constructs (PTPσ Ig1-3_33-327_ and RPTPσ Ig1-3meB_33-331_) were cloned into *Drosophila* pRMHA3 expression vector (Bunch *et al*., 1988). The cDNAs for SALM3 and SALM5 were obtained from ImaGenes GmbH, SALM1 and RPTPσ cDNAs were a kind gift from Dr. Juha Kuja-Panula. The expression constructs included a CD33 signal sequence at the N-terminus of the insert and a C-terminal Fc tag with a preceding the Prescission protease cleavage site. The oligonucleotide primer pairs used for cloning are listed in the Table S1.

The SALM and RPTPσ protein constructs were expressed from stably transfected *Drosophila* S2 cells. Expression was verified by transient transfection using western blot method with goat polyclonal anti-human IgG horseradish peroxidase (HRP) conjugated antibody (Abcam ab98567). All constructs except SALM5 LRR-Ig-Fn_18-505_ were successfully expressed, and were further produced from stable cell lines. For expression from stable cell line of S2 cells, 1.25 ×10^6^ cells per well were plated on a six-well plate at room temperature. After 24 hours, the cells were transfected with 4 μg of DNA containing 1:20 part of selection plasmid pCoHygro. The DNA was diluted into 400 μL of the medium; 8 μL of TransIT insect reagent (Mirus Bio LLC) was mixed with the DNA, and the mixture was incubated for 20 min and added to the cells. After 3 days, the selection was started; the cells and medium were centrifuged and the cells were resuspended into medium with 0.3 mg/ml hygromycin and replated into the same wells. The selection was continued for 3 weeks, with media changed every 6 days, in the same cell culture plate. After 3 weeks, the cells were amplified by splitting them first in low split ratios. The cells were amplified every 6 days until the cell viability was above 95%.

For large scale purification from stable cell lines, the S2 cells were divided 1:10 into HyQ-SFX (ThermoFisher) medium supplemented with 0.15 mg/ml hygromycin, grown in shaker at 25 °C for 1 day, and induced with 0.7 mM CuSO4, and expression was conducted for further 6 days, after which the medium was harvested and cells were pelleted by centrifugation at 7000 rpm for 20 min at 4 °C. The protein was purified using the C-terminal Fc fusion tag with protein-A sepharose (Invitrogen). Samples were eluted with 0.1 M glycine (pH 3.0) directly to neutralizing buffer, 60 mM Tris (pH 7.4) and 300 mM NaCl. The tagged proteins were incubated with Prescission protease for 16 h at 4 °C to remove the Fc tag. Prescission protease was produced as a GST fusion in *Escherichia coli* BL21 (DE3) using plasmid construct developed in pGEX-6P-1 vector (Addgene), and affinity purified with glutathione sepharose (Macherey-Nagel). Cleaved Fc-fusion proteins were passed through a protein-A column, and flow-through containing the cleaved SALM or RPTPσ was collected and gel filtered with Superdex 200 10/300 (GE Healthcare) in 60 mM Tris (pH 7.5) and 300 mM NaCl and concentrated, flash frozen in liquid nitrogen and stored at -80 °C for further use.

### 2.3. Binding affinity measurements and biophysical characterization

Surface plasmon resonance (SPR) measurements on the interaction of SALM3 and SALM5 with RPTPσ variants were carried out using the Biacore T100 system (GE Healthcare) at 25 ^o^C. Fc-tagged SALM3 LRR-Ig-Fn (25 μg/ml) and SALM5 LRR-Ig (25 μg/ml) proteins were immobilized on to a CM5 chip (GE Healthcare) with amide coupling chemistry according to manufacturer’s instructions, and using the running buffer containing 20 mM HEPES (pH 7.5), 150 mM NaCl and 0.005% Tween-20. Two ligands, the RPTPσ Ig1-3 and RPTPσ Ig1-3 meB, were tested for binding against SALM-Fc fusion proteins. The concentrations of wild type RPTPσ and RPTPσ-meB ranged from 0.02 μM to 50 μM.

The size exclusion chromatography-coupled multi-angle static laser light scattering (SEC-MALLS) was used for the characterization of the oligomerization, and monodispersity of the SALMs and RPTP variants. The measurements were done at 0.5 ml/min over an S-200 Superdex 10/300 column (GE Healtcare) in 20 mM Tris (pH 7.4) and 150 mM NaCl with a HPLC system (Shimadzu) and a MiniDAWN TREOS light scattering detector, and Optilab rEX refractive index detector (Wyatt Technology Corp.). Data were then analyzed with ASTRA 6 software (Wyatt Technology Corp.). Proteins were analyzed at 1 mg/ml in 50-100 l volume, expect for SALM3:RPTPσ meB complex, which was measured at molar ratio of 60:80 μM.

### 2.4. Crystallization and structure determination and refinement

SALM5 LRR-Ig construct was concentrated to 8.5 mg/ml and exchanged to 20 mM Tris pH 7.4 and 100 mM NaCl for crystallization. Initial crystals appeared from 0.1 M sodium citrate pH 5.5, 20% w/v PEG 1500 at +20 °C, and were further optimized to 0.1 M sodium citrate pH 5.0, 20% w/v PEG 1500, 0.01 M CuCl_2_ at +4 °C. Crystals were harvested with addition of 10% ethylene glycol or glycerol and flash frozen for data collection. The crystal diffracted to 3.0 Å at best and crystallized in space group F4_1_32, with one monomer in the asymmetric unit (Table 1).

**Table 1.** Crystallographic data collection and refinement

|  |  |
| --- | --- |
| Beam line | ID30B ESRF |
| $\lambda$ (Å) | 0.9677 |
| Temperature (K) | 100 |
| Space group | F4<sub>1</sub>32 |
| *Unit cell* |  |
| a = b = c (Å) | 254.0 |
| Resolution (Å) | 50-3.0 |
| Observations | 159812 |
| Unique | 14488 |
| Rmerge (%) | 13.6 (222.6) |
| CC(1/2) | 98.30 (40.90) |
| I/(I)$\sigma$ | 10.59 (1.10) |
| *Refinement* |  |
| Resolution (Å) | 30-3.0 |
| Rwork / Rfree (%) | 25.20 / 28.20 |
| No. Reflections / Rfree-set reflections | 14438 / 721 |
| Average B-factor (Å<sup>2</sup>) | 142.1 |
| LRR-domain (Å<sup>2</sup>) | 126.1 |
| Ig-domain (Å<sup>2</sup>) | 200.2 |
| No. of atoms (protein) | 2457 |
| r.m.s.d bond lengths (Å) | 0.01 |
| r.m.s.d bond angles (°) | 1.36 |
| Ramachandran plot<br>(favored/outliers) (%) | 91.60 / 1.24 |

The SALM5 crystal structure was solved using the BALBES automated molecular replacement pipeline with CCP4 (https://www.ccp4.ac.uk/ccp4online/) (Long *et al*., 2008). Molecular replacement via the BALBES pipeline was able to find an initial solution for the LRR-domain of SALM5, using the LRR domain of AMIGO-1 as the initial template (PDB:2XOT) (Kajander *et al*., 2011) as a template, but no solution for the Ig-domain. After molecular replacement and initial refinement with REFMAC (Murshudov *et al*., 2011), the R-factors for the initial LRR domain model were R_work_/R_free_ = 43.2%/45.9%. Subsequently several runs of the autobuild function in phenix software suite (Terwilliger *et al*., 2008) and “morphing” (Terwilliger *et al*., 2013) were used to initially build the model from scratch using the molecular replacement solution to correct the model and trace partially the Ig-domain during the refinement. Finally the Ig-domain was modeled using Swiss-modeller (Bordoli *et al*., 2008), and the best minimal model was aligned with the partial crystal structure model built in the 2Fo-Fc electron density traceable for the Ig-domain and rerefined with the autobuild “rebuild-in-place” and “morphing” options in phenix. All of this was done with several iterative cycles of model building with Coot (Emsley and Cowtan, 2004) with most of the surface side chains and the loop between Ile328 to Leu335 removed during building due to lack of density. This way a model for the Ig-domain was completed and final rounds of refinement were done with BUSTER (Bricogne *et al*., 2017). Final R-factors were R_work_/R_free_ = 25.2%/28.2% and model had geometry restrained to standard expected quality (Table 1).

### 2.4. Small-angle X-ray scattering data collection and analysis

Small-angle X-ray scattering (SAXS) data on the proteins were measured for SALM5 in batch mode at Diamond Lightsource on the B21 beam line, and for SALM3 at the ESRF beam line BM29 (Table S2). SALM5 sample was measured in 30 mM Tris-Cl pH 7.5, 150 mM NaCl, 3% glycerol at 1, 3 and 5 mg/ml in 50 μl volume. The SALM3 samples were measured by SEC-SAXS. Sample compartment and exposure cell were cooled to 4 °C (Table 3). An SD200 10/300 GL (GE Healthcare) column was used at 0.5 ml/min in 20 mM Tris HCl pH 7.5, 100 mM NaCl, 0.02% NaN_3_. For the SALM3 LRR-Ig construct 45 μl of sample at 8.8 mg/ml, was used for SEC-SAXS analysis and for SALM3 LRR-Ig-Fn 47 μl of 15.6 mg/ml sample. Data processing was performed automatically using the EDNA online data analysis pipeline using tools from the ATSAS 2.5.1 (Incardona *et al*., 2009) generating radially integrated, calibrated, and normalized one-dimensional profiles for each frame. DATASW (Shkumatov and Strelkov, 2015) was used for calculation of the invariants (I(0), Rg, and molecular weights (Mw).

For SALM3 constructs, an elution profile was generated with the I(0)/Rg variation plotted versus recorded frame number. The averaged data, corresponding to frames where Rg is stable and shows linearity, were further processed using ATSAS package (Petoukhov *et al*., 2012). PRIMUS software from the ATSAS software suite version 2.8.0 was used for primary data processing (Franke *et al*., 2017). Parameters for each sample are given in Table S2. Glycans were added to the models with the GlyProt server (www.glycosciences.de/modeling/glyprot/php/main.php).

Ensemble Optimization Method (EOM) from ATSAS package was used with default parameters (‘Random-coil’ chain type) resulting in a random pool of 10 000 C-α trace models. Next, a computational pipeline FULCHER (Shkumatov A. et al, unpublished data) was used to convert C-alpha trace models to all-atom models, with subsequent model validation using the MOLPROBITY clash score of 40. Finally, the genetic algorithm GAJOE was run 10 times to obtain an ensemble of models that best describes the experimental SAXS data.

## 3. Results and Discussion

### 3.1. Overall Structure of SALM5 reavels a dimeric assembly

We have solved the structure of SALM5 at 3.0 Å resolution. The protein crystallized in the cubic space group F4_1_32 (see Table 1). The structure reveals a typical extracellular LRR-domain with the LRRNT and LRRCT capping subdomains with two stabilizing disulphides each, and seven LRR repeat between these. Based on sequence data it has been unclear whether there was six or seven LRR repeats, the structure confirms the presence of seven LRR repeats. Overall, the extracellular LRR domains tend to be highly variable in size, e.g. the neuronal LRRTMs have 10 LRR-repeats (Lauren *et al*., 2003), while others have less, e.g. Slit LRR domains have 4-6 repeats (Rothberg *et al*., 1990), while Toll-like receptors have 19-25 repeats (Botos *et al*., 2011) and several proteins have consecutive separate LRR-domains in tandem, such as SLITRKs and Slits (Rothberg *et al*., 1990; Beaubien and Cloutier, 2009). While all eukaryotic extracellular LRR domains share the disulphide-linked capping domains at the N- and C-termini of the LRR-domains. Also, combinations of LRR and Ig-domains and LRR and Fn-domains, such as in the SALM-family proteins, are common among the neuronal cell surface LRR-proteins, which typically reside on the post-synaptic membrane.

In SALM5, the N-terminal LRRNT capping subdomain (residues Phe17-Arg52) has the structure typical of an extracellular eukaryotic N-terminal capping motif (Park *et al*., 2008). The LRRNT β-hairpin at the N-terminus of the LRR domain β-sheet structure (Fig. 1) has slightly longer β-strands than the rest of the LRR β-sheet. The C-terminal LRRCT-capping subdomain (Gln216-Cys284) is quite divergent from the typical LRRCT structure. It starts with an unusual large insertion, disordered in the crystal structure (Gln216-Phe233) before the last β-strand of the extended LRR-domain β-sheet; the first ordered residue is Ala234 (marked in Figure 1). The rest of the LRRCT consists of only one regular α-helix and several turns with two disulphide bridges stabilizing it. The capping LRRCT appears to be more loosely packed against the LRR repeats than in related proteins, with somewhat irregular secondary structure. The looser structure is possible because of the large insertion in the LRRCT. As is typical, the LRR domain ends at the last Cys-residue of the second LRRCT disulphide (Cys285-Cys246). The other disulphide bridge in the LRRCT is formed by the Cys244 and Cys263 residues.

**Figure 1.**
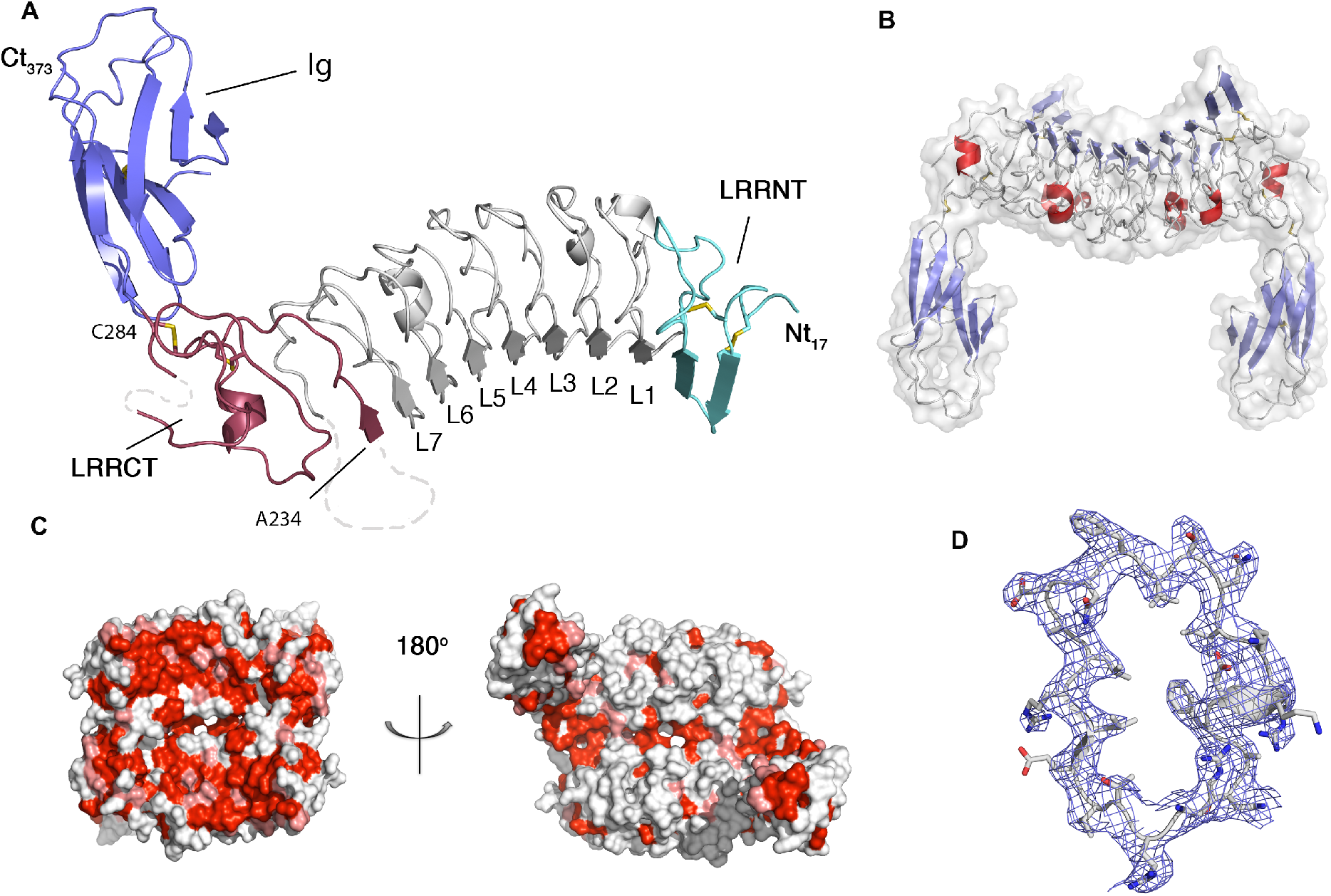
The SALM5 crystal structure and the dimer assembly of the LRR-Ig fragment. A) The monomeric SALM5 LRR-Ig construct structure. Disulphide bridges drawn as sticks and missing parts for the LRR domain shown as dashed lines, Ig-domain is indicated with “Ig” and LRRNT (cyan) and LRRCT (dark red) capping domains are indicated as “LRRNT” and “LRRCT”. Also the first ordered residue of the LRRCT, Ala234, and the last residue of the whole LRR domain, Cys284, are marked in the figure. The N- and C-termini of the protein are marked with “Nt” and “Ct” and the residue number. LRRs 1-7 are labeled as L1 to L7. B) Dimeric assembly observed in the crystal structure as a side view with transparent surface, coloured by secondary structure (β-strands as blue, helices in red). C) Amino acid conservation plotted on the surface indicate the top-concave LRR dimer surface (left) as the most conserved area. Red and light red indicate the most conserved residues as calculated by CONSURF (Ashkenazy, Abadi et al. 2016) for SALM3 and SALM5. Convex side view (right) shows little conservation in the LRR domain or the Ig-domain. D) A 2Fo-Fc electron density contoured at 1 σ for a representative part of the LRR domain for the residues of the first LRR repeat (L1 in A). All molecular images were made with PyMoL.

In the mouse SALM5, LRR domain extra electron density can be seen in the predicted Asn73 N-glycosylation site, but the sugar moiety could not be modeled reliably. The other two predicted N-glycans reside in the Ig-domain at residues Asn330 and Asn339. These are disordered in the crystal structure. Overall, based on their location it would appear that the glycans do not have functional significance.

The SALM5 Ig-domain (Glu285-His373) follows immediately after the LRRCT-capping subdomain (Fig 1). The Ig-domain appears fairly disordered overall and apparently somewhat mobile in the crystal as it has poor electron density with higher B-factors compared to the LRR domain (Table 1). Nevertheless it was possible to trace its position up to the last residue of the fold, His373, based on the initial partial model (see methods). Some parts of the Ig-domain are weak in the density and loop between two β-sheets of the Immunoglobulin β-sandwich fold at Ile328 and Leu335 is disordered. A part of typical electron density for the LRR domain is shown in Figure 1, covering the region for the first LRR repeat.

The refined crystal structure forms a dimer with C2 symmetry with a subunit interface formed by the LRR-domains edge-to-edge, the top surfaces of the LRR β-sheets packed against each other (Fig. 1), with a surface area of 1046 A, which is clearly the largest intermolecular interface found in the crystal, as measured with the PISA-server (Krissinel and Henrick, 2007). Interestingly though, the Ig-domains form trimeric assemblies in the crystal, which might be able to further contribute to possible clustering of the molecules on the membrane. In the full length protein the Ig-domain is followed by a linker region of ca. 40 residues (Ile374-Thr413) and a fibronectin type-III domain on the extracellular side before the transmembrane helix and cytosolic tail. The dimeric LRR-Ig structure has dimensions of *ca.* 80 Å x 65 Å x 55 Å.

The LRR domains in the dimer form a shallow bowl-shaped surface towards the synaptic cleft formed (Fig. 1) by the two LRR domain concave surfaces, while the Ig-domains point “downwards” at almost 90 degree angle towards the cell-membrane from the putative LRR-dimer concave ligand binding platform, which shows high degree of conservation between the known RPTP binding SALM3 and SALM5 proteins (Fig. 1). In total we find 31 highly conserved residues on the concave surface on the LRR-domain beta-sheet from LRRNT to the seven LRR repeat (Table 2). In practice, the whole surface appears highly conserved (Fig. 1), while the overall sequence identity between the SALM3 and SALM5 proteins is ca. 50-60% and the convex side of the LRRs (or the Ig-domain) do not show significant continuous areas of conservation (Fig. 1).

**Table 2.** Most conserved residues on the concave SALM5 LRR surface.

| **LRR region** | **Conserved residues<sup>§</sup>** |
| --- | --- |
| LRRNT | Asn32 *, Ala34 *, Leu36 |
| LRR1 | Val54, Glu55, Arg57, Asp60 |
| LRR2 | Val78, Asp79, Thr81, Ser83, Arg84 |
| LRR3 | Arg102, Ala103*, His105 |
| LRR4 | His126*, His127, Ile129, Asn131 |
| LRR5 | Glu150, Glu151*, Asp153, Ser155 |
| LRR6 | His174, Thr175*, Ser177*, Asp179 |
| LRR7 | Arg199, Asp201, Thr203, Ser204 |

<sup>§</sup>Residues with Consurf-server scores: 9\* and 10.

**Table 3.** The binding affinities (K_d_-values) of SALM3 LRR-Ig-FN and SALM5 LRR-Ig for RPTPRσ and RPTPRσ-

| | PTPσ | PTPσ meB |
| --- | --- | --- |
| SALM3 LRR-Ig-FN | $23.3 \pm 2.0$ μM | $2.2 \pm 0.2$ μM |
| SALM5 LRR-Ig | $10.6 \pm 1.8$ μM | $5.4 \pm 0.7$ μM |

The dimer interface is formed by hydrophobic interactions at the N- and C-termini on the LRR domain (Fig. 2) involving residues Phe62 and Leu43 on one side and Leu260, Ala264 and Thr262 on the other side, partly packing against the peptide backbone turns of the opposing domain, and by hydrogen bonding interactions in the middle of the interface by side chains of Arg110 and Asn158 from opposing monomers, Arg110 also hydrogen bonds to the backbone carbonyl of Asn157. Also Gln134 is possibly involved in the hydrogen-bonding network (Fig. 2) but its position could not be accurately modeled in the 3.0 Å resolution 2Fo-Fc electron density map.

**Figure 2.**
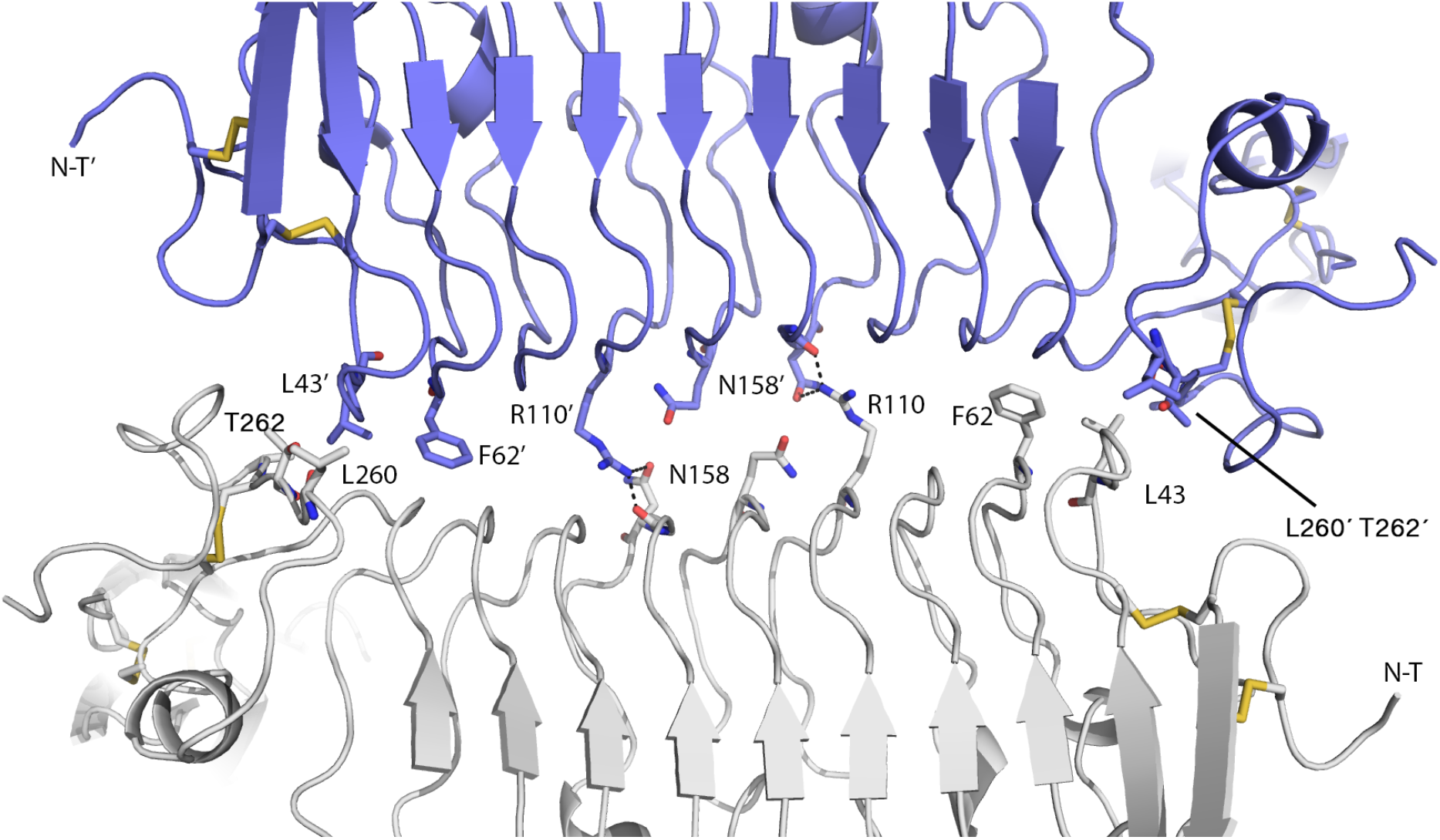
The SALM5 dimer interface formed by the LRR-domains. Residues involved in dimer interface contacts are labeled, for further details see text. In the middle of the interface a conserved set of polar residues contribute to hydrogen bonding between the monomers, towards the N- and C-termini of the monomers at the dimer interface more hydrophobic interactions contribute to the interface formation. Monomers are shown in light grey and blue, N-T = N-terminus. Ig-domain is hidden behind the plane of the paper.

### 3.2. Comparison to other LRR proteins

Based on a DALI-search (Holm *et al*., 2008) we found several LRR domain proteins with high Z-scores. The highest Z-scores were found for the LRR domain for the Hagfish variable lymphocyte receptor (VLR) (Z-score 27.6) domain with seven LRRs (PDB:2O6Q) and the platelet glycoprotein Ib alpha (GP1ba; PDB:1M0Z) (Z-score 25.9), also with seven LRRs. The GP1ba LRR domain aligns with an r.m.s.d. of 1.462 Å with the SALM5 LRR domain (Fig 4). Interestingly, both SALM5 and GP1ba have a large loop insertion in the LRRCT subdomain, which in the GP1ba wraps around the von Willebrand factor ligand, structuring into a β-hairpin structure on ligand binding (Huizinga *et al*., 2002). Similarly, the β-sheet of LRRNT is extended in both, and in GPIba bent towards the binding site on the concave surface of its LRR domain inducing larger curvature, and wrapping around the ligand. It can be hypothesized that the long insertion loop in SALM5 also might have a functional role in recognition of the ligand, since the insertion at this position is also present in the whole SALM protein family.

Also the SLTRK 1st LRR domain, with six LRRs, aligns well (r.m.s.d. 1.429 Å) with our SALM5 structure – this is notable as both recognize the same RPTP-ligands (see below). Previously we have characterized the structures of the neuronal AMIGO family proteins (Kajander *et al*., 2011), which have six LRR followed by an Ig-domain, but with a different type of dimer arrangement, where the LRR concave surfaces form the dimer interface. The SALM LRR domain dimer appears unique in the known cell surface LRR-protein proteins by forming a continuous double β-sheet surface by the two LRR domains (Fig. 1).

### 3.3. Biophysical characterization of SALM family proteins

Solution measurements by SEC-MALLS and SAXS analysis confirm the dimeric nature of SALM-family proteins in solution, as dimers were observed for all SALM1_LRR-Ig-FN_, SALM3_LRR-Ig_, SALM3_LRR-Ig-FN_ and SALM5_LRR-Ig_ by SEC-MALLS (Fig. 3), with observed molecular weights (Mw) of 80.7 and 83.2 kDa for SALM3 and SALM5_LRR-Ig_ constructs, and 121 kDa for the Fn-domain containing constructs. These match well with expected theoretical dimer Mws based on the calculated monomer Mws of 39.8 and 39.1 kDa for the short constructs, as well as the longer ectodomain constructs, which have theoretical Mws of 52 and 54 kDa for a monomer. All theoretical values are without N-glycans, which contribute ca. 3-4 kDa per protein monomer, due to the paucimannose-type glycans present in insect cell-produced proteins (Shi and Jarvis, 2007).

**Figure 3.**
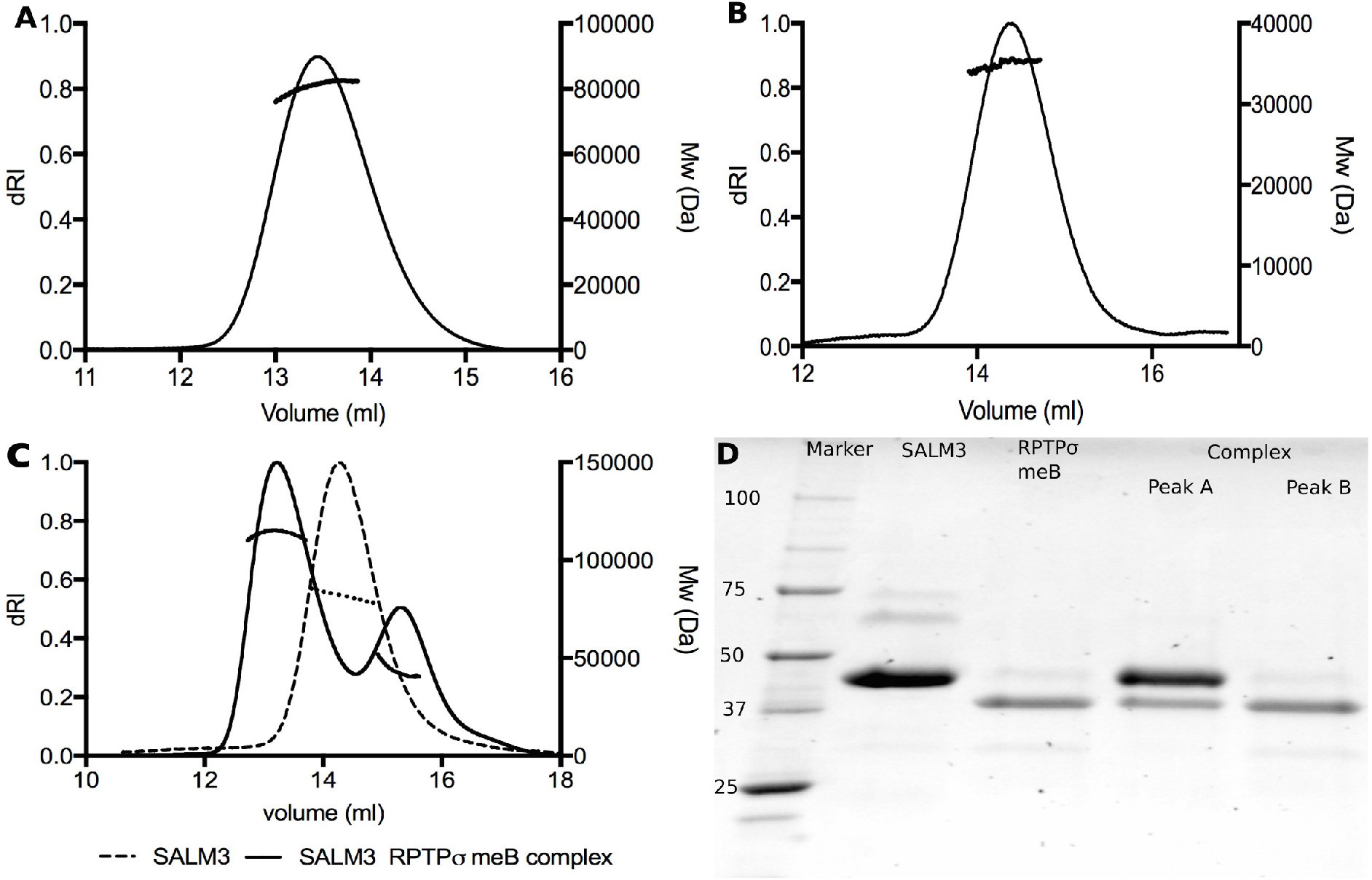
Characterization of the SALM proteins oligomerization and ligand binding by analytical size exclusion chromatopgrahy and static light scattering. A) SALM5 LRR-Ig domain construct B) The RPTPR Ig1-3-mEB construct. C) SALM3 LRR-Ig:RPTP-meB complex (solid line) and SALM3 LRR-Ig construct alone (dashed line). D) An SDS-PAGE analysis of the first (“A”) and second (“B”) peaks of the SALM3:RPTP-meB complex, verifying the presence of both proteins in the complex peak (“A”). Calculated Mws (right Y-axis) are plotted over the protein (dRI signal) peaks with a solid or dashed line.

**Figure 4.**
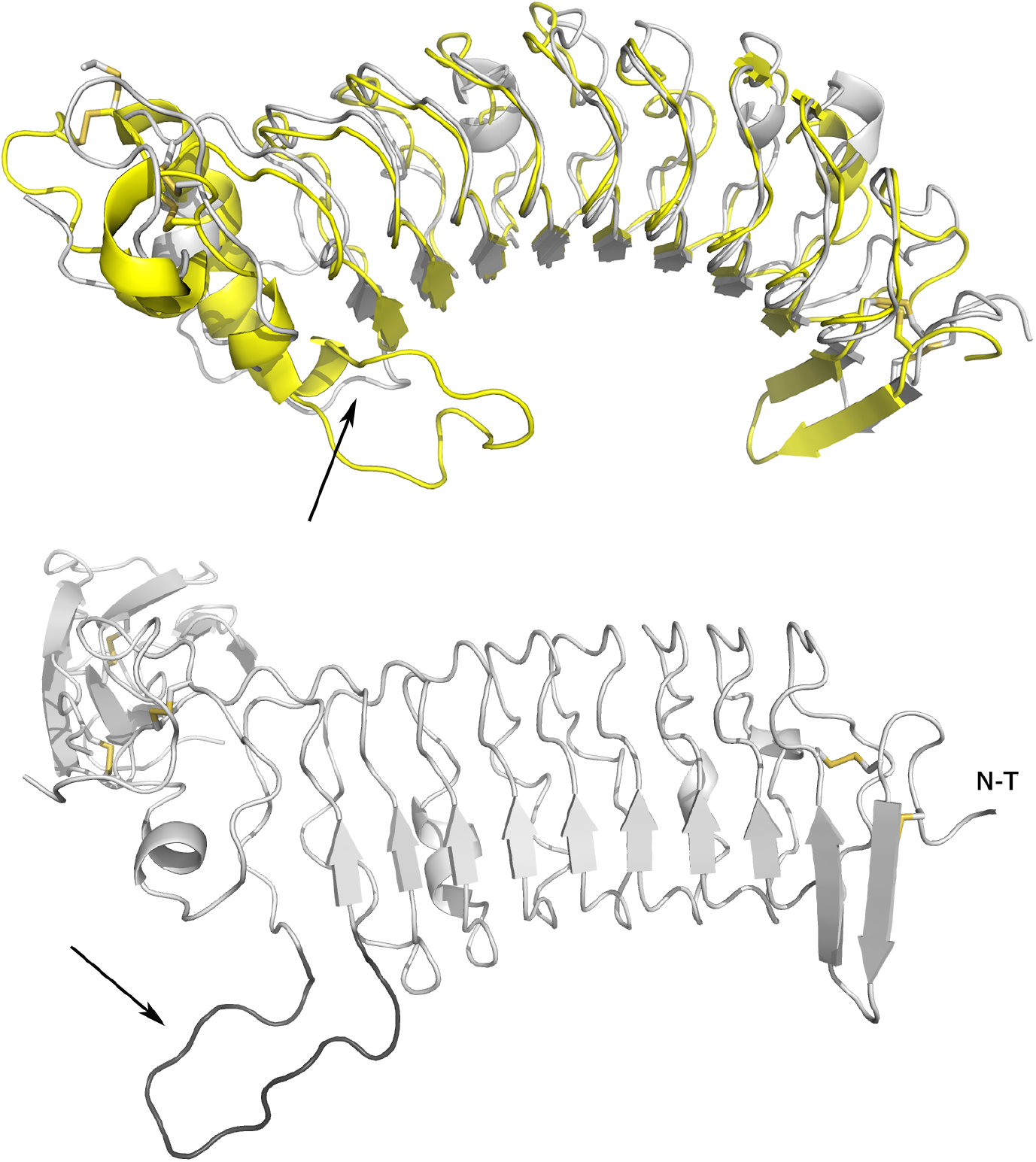
Superimposition of SALM5 with LRR domain of platelet Glycoprotein Ib alpha (GPIba). Upper figure: The alignment of the seven repeat LRR domains (GPIba in yellow and SALM5 in grey). Lower figure: the crystal structure of the SALM5 LRR-Ig fragment with the disordered loop Gln216-Phe233 modelled (in dark grey). The arrow indicates the loop insertion at the start of the LRRCT capping subdomain in SALM5, while the insert in GPIba is in the middle of the LRRCT. N-T; N-terminus. Structures were aligned and molecular images made with PyMoL.

To further confirm the oligomeric state of SALM-proteins in solution, SAXS experiments were performed (Fig. 5, Table S2). The scattering profiles show few features for SALM3 and SALM5 LRR-Ig constructs and almost a featureless curve in case of the whole SALM3 ectodomain construct (SALM3 LRR-Ig-Fn) (Fig. S1, Fig. 5). A dimensionless Kratky (dKratky) plot (Fig. 5) reveals two peaks for SALM3 LRR-Ig, whereas the second peak was less pronounced for SALM5 LRR-Ig and almost absent for the SALM3 LRR-Ig-Fn construct, indicating that ectodomain construct is the most flexible among three. Similarly, all P(r) functions present a single peak with a smooth and extended P(r) distribution function, particularly in case of the ectodomain construct (Fig. 5). Molecular weight estimation from SAXS curves of SALM3 and SALM5 LRR-Ig constructs indicated values consistent with a dimer model, whereas for SALM3 LRR-Ig-Fn increased Mw estimates can be due to presence of the long disordered linker region between the Ig- and the Fn-domains (Table S2, Fig 5B).

**Figure 5.**
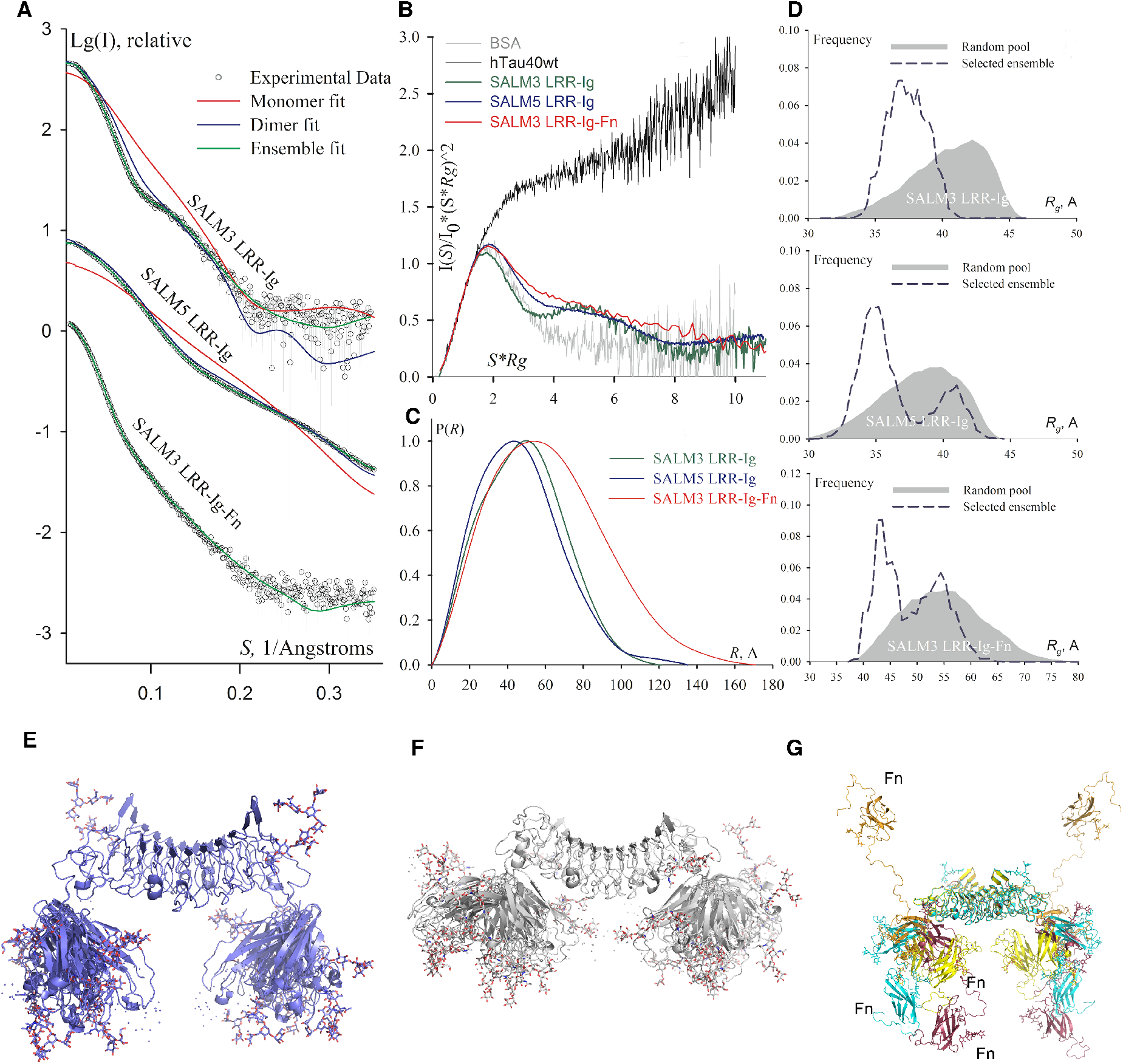
Solution structures of SALM5 and SALM3 support a conserved dimer interface and assembly. Small angle X-ray scattering analysis of SALM3 and SALM5 constructs. A) Scattering intensity profiles with corresponding fits to a monomer, dimer and ensemble model. B) Dimensionless Kratky plots in comparison with globular BSA (gray line) and natively unfolded human hTau40 protein (black line). C) Distance distribution function, P(R), estimated for the studied constructs using GNOM. D) Flexibility assessment: *Rg* distribution for the random pool of models (gray) and for the selected ensemble of conformations that fit the scattering intensity curve (dashed dark blue). E-G) The ensemble models for dimers of SALM3 LRR-Ig (E, blue), SALM5 LRR-Ig (F, grey) and SALM3 LRR-Ig-Fn constructs (G, multiple colours). Dimers are viewed from the side as in Fig 1b. It can be seen that the Ig-domains form flexible ensembles around the fixed LRR-domain dimer. SALM3 LRR-Ig-Fn construct shows significantly increased flexibility, probably due to the linker between the Ig- and Fn-domains. Positions of the Fn-domains in the selected ensemble models are marked for one monomer (in G). Glycans are shown as sticks.

For validation of obtained SALM5 crystal structure, theoretical scattering profiles were calculated for monomer and dimer model of SALM5 and compared to the experimental scattering profile. The resulting fits unambiguously indicated that both LRR-Ig scattering curves resemble dimer state in the crystal (Fig. 5). To reconstitute the scattering curves, missing flexible termini were modelled and data described using an ensemble approach (Bernado *et al*., 2007). The obtained fits show excellent agreement to the scattering data (Fig. 5). In case of SALM3_LRR-Ig_, *Rg* distribution of the selected ensemble is shifted to the left, suggesting a rather compact structure of the construct, consistent with observed “bumps” in the scattering curve and the second peak in the dKratky plot (Fig. 5). In case of SALM5_LRR-Ig_ construct, additional smaller peak was observed (Fig. 5), which corresponds to presence of extended confirmations. The whole SALM3 ectodomain construct (SALM3_LRR-Ig-FN_), on the other hand, shows a rather broad *Rg* distribution of the selected ensemble (Fig. 5), indicating higher flexibility as compared to the other two constructs.

The conserved structural arrangement between the homologous SALM proteins, as well as the sequence conservation (Fig. 1), suggests a conserved mechanism of action between at least the SALM3 and SALM5 proteins for ligand binding, as they both bind the same presynaptic RPTPσ-ligands.

### 3.4. Ligand binding by SALM3 and SALM5 and model for binding site for RPTP-ligands

We determined the binding affinities of SALM3 and SALM5 extracellular domains towards RPTPσ and RPTPσ-meB splice variant by SPR. SALM3 showed a clear preference for RPTPσ-meB splice variant, while for SALM5 the data indicated similar affinities for both (Fig 6.). Overall, SALM3 had slightly higher affinity and was more specific towards the RPTPσ-meB ligand, with 10-fold lower affinity observed by SPR for the RPTPσ lacking the mini-exon B between the Ig2- and Ig3-domains of the RPTPσ ligand. For SALM5 similar affinities were observed with or without the meB-insert of RPTPσ. Both proteins had similar low micromolar range affinities for the meB-containing splice variant ligand (Table 3). Previously, SALM3 has been reported to have a preference for the meB splice variant (Li *et al*., 2015). Notably, we also did not observe any binding of SALM1 to SALM3 or SALM5, contradicting the earlier aggregation results (Nam *et al*., 2011).

**Figure 6.**
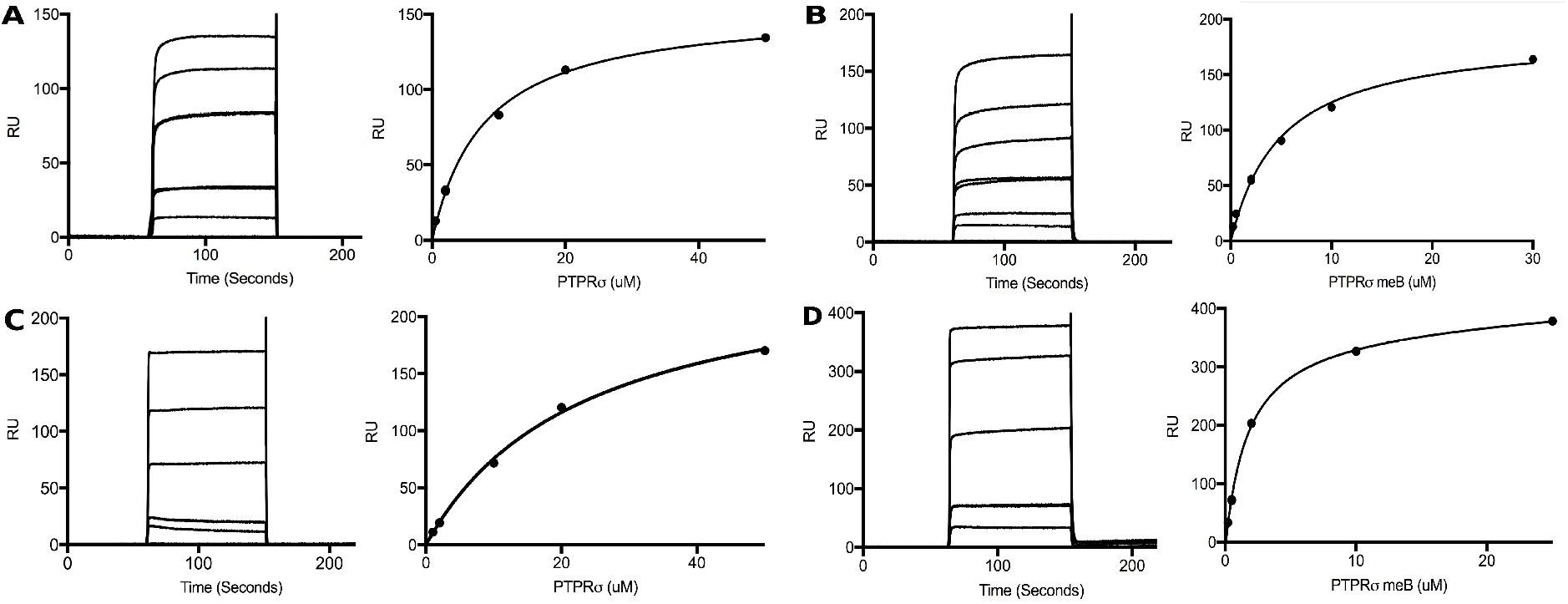
Binding of SALM3 and SALM5 to the RPTP ligand as measured by surface plasmon resonance. Binding of SALMs to RPTPσ/RPTPσ-meB. SALM3 LRR-Ig-Fn and SALM5 LRR-Ig were tested for binding to RPTPσ and RPTPσ-meB Ig1-Ig3 constructs by SPR. A) SPR sensograms (left) and equilibrium binding curve plotted (right) for binding of SALM5 LRR-Ig to RPTPσ. B) SPR sensograms (left) and equilibrium binding curve plotted (right) for binding of SALM5 LRR-Ig to RPTPσ meB. C) SPR sensograms (left) and equilibrium binding curve plotted (right) for binding of SALM3 LRR-Ig-Fn to RPTPσ. D) SPR sensograms (left) and equilibrium binding curve plotted (right) for binding of SALM3 LRR-Ig-Fn to RPTPσ meB.

Based on the earlier reported structures of SLITRK in complex with RPTP Ig-domains (Um *et al*., 2014; Yamagata *et al*., 2015), we decided to compare the two systems. Given the clear sequence conservation pattern among SALM3 and SALM5 proteins when displayed on SALM5 surface (Fig. 1), it seems obvious that SALMs will also recognize their ligand via the unique concave dimeric LRR domain surface, possibly suggesting a new type of ligand recognition for the LRR-adhesion receptors. Overall, LRR domains typically recognize their targets utilizing the concave surface (Helft *et al*., 2011). In order to compare the two systems, we aligned the SALM5 LRR domain and the SLITRK LRR domains from N-terminus onwards and observed how the RPTP ligand would fit on the SALM dimer “top” concave surface (Fig. 7). Obviously no direct fit is expected, but based on the alignment, it can be suggested that the RPTP ligand would not fit very well in a 2:2 complex onto the SALM5 dimer, unless the binding occurs in a very different manner. This would have to involve the non-conserved regions on the LRR domains, away from the dimer interface and the “top”-surface (Fig. 7). We then performed assays to verify the stoichiometry, and based on the SEC-MALLS analysis (Fig 3) of the SALM3-PTPσ Ig1-Ig3 complex, peaks corresponding to 2:1 complex and dissociated RPTPσ ligand were observed (Fig. 3). We therefore suggest that the RPTPσ-ligand is recognized with 2:1 stoichiometry. It still possible that there is a second binding event with lower affinity that we are unable to observe, which would result in some form of 2:2 complex. Nonetheless, it would appear that the 2:1 complex is the major quaternary state for the complex, based on the SEC-MALLS results, excluding possible clustering effects by other factors.

**Figure 7.**
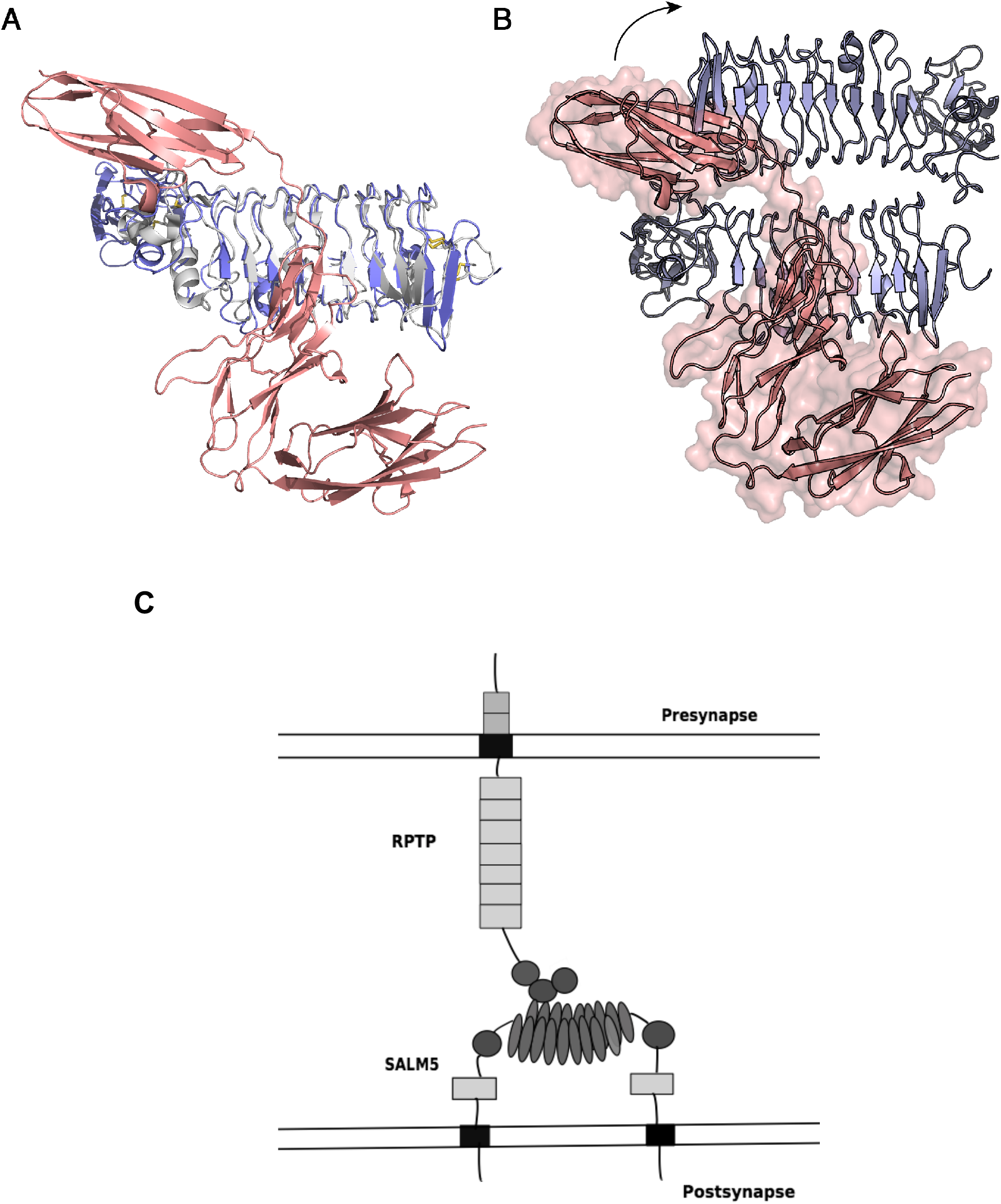
A model for the interaction complex of SALM5 with RPTP ligand and comparison with SLITRK complex. A) The monomeric SLITRK:RPTP complex with the SLITRK and SALM5 LRR domains aligned, see text (SLITRK in grey, SALM5 in blue, RPTP, red). B) Potential interaction model for SALM5 dimer and RPTP ligand. Arrow indicates changes in the orientation of the Ig3-domain of RPTPσ (in red, with transparent surface) required to avoid steric clashes in the hypothetical SALM5:RPTPσ complex. Approximate position of the ligand is consistent with surface conservation in SALM5 and SALM3 proteins (See Fig. 1). C) Schematic 2:1 interaction model at the synapse for SALM5 interaction with presynaptic monomeric RPTP. Rectangles = Fn-domains, spheres = Ig-domains, ellipses the SALM LRR repeats. Further clustering of the complex could occur via other interactions by either partner.

## 4. Conclusions

We show that SALM5 appears to function as a dimeric synaptic organizer molecule – the crystal structure revealing a bowl-shaped top surface formed by the dimerized LRR-domains. We find that SALM1 and SALM3 ectodomains also, and therefore most likely the whole protein family, form dimeric structures, presumably via their LRR-domains. Our SAXS analysis confirms that SALM3 LRR-Ig fragment has a very similar structure to the SALM5 crystal structure. Both SALM3 and SALM5 in our studies are able to bind the RPTP ligands with meB splice site with similar affinity while SALM3 appears more selective. Based on our results, we suggest a 2:1 stoichiometry for RPTP-ligand binding, and locate the ligand binding site at the extended concave LRR “double “-sheet” surface of the dimeric SALM ectodomain based on the sequence conservation data and the structural organization.

## Acknowledgements

We thank the staff at Diamond Lightsource and ESRF beamlines for help with data collection on beam lines BM29 (ESRF), Diamond B21, and ESRF ID30B. Crystals were initially screened at the Biocenter Finland crystallization core facility at Institute of Biotechnology, Helsinki, Finland. We also wish to thank Dr. Juha Kuja-Panula for the gift of RPTP and SALM1 cDNAs, Dr. Katja Rosti for work in the initial phases of the project on SALM1 and SALM5 and Seija Mäki for help with crystallization. The coordinates and structure factors have been deposited in the protein data bank with accession code 6F2O.

